# Locomotor response to acute stressors requires hypothalamic-pituitary-interrenal axis activation and glucocorticoid receptor

**DOI:** 10.1101/291914

**Authors:** Han B. Lee, Tanya L. Schwab, Ashley N. Sigafoos, Jennifer L. Gauerke, Randall G. Krug, MaKayla R. Serres, Dakota C. Jacobs, Ryan P. Cotter, Biswadeep Das, Morgan O. Petersen, Camden L. Daby, Rhianna M. Urban, Bethany C. Berry, Karl J. Clark

## Abstract

When vertebrates face acute stressors, their bodies rapidly undergo a repertoire of physiological and behavioral adaptations, which is termed the stress response (SR). Rapid physiological changes in heart rate and blood sugar levels occur via the interaction of glucocorticoids and their cognate receptors following hypothalamic-pituitary-adrenal (HPA) axis activation. These physiological changes are observed within minutes of encountering a stressor and the rapid time domain rules out genomic responses that require gene expression changes. Although behavioral changes corresponding to physiological changes are commonly observed, it is not clearly understood to what extent HPA axis activation dictates adaptive behavior. We hypothesized that rapid locomotor response to acute stressors in zebrafish requires HPI axis activation. In teleost fish, interrenal cells (I) are functionally homologous to the adrenal gland cortical layer. We derived 8 frameshift mutants in genes involved in HPI axis function: two mutants in exon 2 of *mc2r* (adrenocorticotropic hormone receptor), two in each of exon 2 and exon 5 of *nr3c1* (glucocorticoid receptor), and two in exon 2 of *nr3c2* (mineralocorticoid receptor). Exposing larval zebrafish to mild environmental stressors, acute changes in salinity or light illumination, results in a rapid locomotor response. We show here that this locomotor response requires a functioning HPI axis via the action of *mc2r* (adrenocorticotropic hormone receptor) and the canonical glucocorticoid receptor encoded by *nr3c1* gene, but not mineralocorticoid receptor (*nr3c2*). Our rapid behavioral assay paradigm based on HPI axis biology may prove useful to screen for genetic, pharmacological, or environmental modifiers of the HPA axis.

**Significance:** Altered HPA axis activity is acknowledged as a causative and critical prognostic factor in many psychiatric disorders including depression. Nonetheless, genome wide association studies (GWAS) on depression have revealed conflicting findings about susceptibility loci, while identifying several genetic loci that warrant further investigations in the process. Such findings indicate that psychiatric disorders with complex genetic foundations require functional studies as well as genetic analyses. We developed a sensitive behavioral assay paradigm that leverages the genetic amenability and rapid development of zebrafish and demonstrated that our assay system reliably detects changes in HPA axis responsiveness. Our functional genetics and behavioral assay approach provides a useful platform to discover novel genetic, pharmacological, or environmental modifiers of the HPA axis.

## Introduction

The stress response (SR) is an organism’s response to perceived threats to homeostasis (1, 2). Intense acute stress or prolonged stress that overwhelms the body’s SR system is detrimental to an organism’s health (3). Such stress beyond an organism’s coping capacity is associated with the onset or aggravation of a broad spectrum of psychiatric disorders such as major depressive disorder, anxiety disorders, post-traumatic stress disorder, and substance use disorders (4–7). The SR is mediated primarily by the hypothalamic-pituitary-adrenocortical (HPA) axis. Not only are alterations in HPA axis activity one of the most consistent findings among people with psychiatric disorders, but also normalization of HPA axis activity is a critical parameter that determines patients’ prognoses and risk of relapse (8–12). To devise effective therapeutic strategies for complex psychiatric disorders, it is essential to advance our understanding regarding the pathways and genes that regulate the HPA axis and, in turn, how alterations in HPA axis activity may lead to psychiatric illness.

Evolved and conserved in vertebrates, activation of the HPA axis leads to the secretion of glucocorticoids (GCs) from the adrenal gland cortical layer in tetrapods (13, 14) and from the interrenal cells (HPI axis) in teleost fish (15). GCs, such as cortisol in humans and zebrafish or corticosterone in rodents, are effector molecules that modulate the SR (16–18). GC signaling via cognate receptors (corticosteroid receptors) mediates pleiotropic effects of SR in a tissue-specific manner. Corticosteroid receptors (CRs) include type I (mineralocorticoid receptor (MR) encoded by *nr3c2* (nuclear receptor subfamily 3 group c member 2)) and type II (glucocorticoid receptor (GR) encoded by *nr3c1*) receptors. There is a potentially yet-to-be-identified group of membrane-associated G-protein coupled receptors (GPCRs) (19–21). As members of nuclear receptor family transcription factors, GR and MR have been most extensively investigated as agents of gene expression changes. However, the temporal characteristics of GC activity point to biphasic actions of GC-CR interactions. Slower responses involving changes in gene expression usually take more than 30 min to manifest and are known as the genomic response. Rapid nongenomic responses occur within only a few minutes and involve various downstream signaling pathways (22–24). Mounting evidence suggests that nuclear receptor family transcription factors (GR and MR), as well as putative membrane-associated receptors, play a role in rapid nongenomic GC signaling that is transcription-independent (25–27).

It is largely unknown how and to what extent such rapid GC signaling regulates HPA axis activity. We have a limited understanding of what factors modulate rapid GC signaling, the role of rapid GC signaling in HPA axis regulation, or the effects a dysregulated HPA axis has on behaviors due to changes in rapid GC signaling. Our group previously reported that larval zebrafish respond to hyperosmotic stress (application of sodium chloride) with increased frequencies of locomotion (28). The zebrafish, a teleost with conserved cortisol-synthetic pathways and SR genes, serves as an effective model system for genetic and pharmacological manipulations to investigate the SR. We aimed to develop a sensitive behavioral assay paradigm that captures alterations in HPA axis activity with a quantifiable readout in order to establish a causal relationship between perturbed HPA axis activity and altered locomotion. Here we demonstrate, using larval zebrafish, that rapid locomotor response to acute stressors requires cortisol secretion via the action of melanocortin receptor type 2 (*mc2r*; adrenocorticotropic hormone (ACTH) receptor) and depends on the canonical corticosteroid receptor type II (glucocorticoid receptor) encoded by *nr3c1*. Since such locomotor changes occur within minutes of stressor applications, our assay system provides an effective platform to screen for genetic, pharmacological, or environmental modifiers of rapid responses of the HPA axis and will contribute to a better understanding of the role that rapid GC signaling plays in shaping HPA axis activity and the SR at the cellular and organismal levels.

## Results

### Conception of an acute stress assay paradigm

We set out to establish an acute stress assay behavioral suite. The stress assay suite comprises two stress paradigms and a noxious stimulant locomotor control paradigm. We designed acute stressors that capitalize on zebrafish biology, primarily that they are diurnal, freshwater fish. One stressor includes abrupt changes in light and the other includes an acute increase in salinity. A noxious stimulant, cinnamon oil, is used as a positive control for locomotor activity (Fig. 1). Behavioral assays are performed on 5 days-post-fertilization (dpf) since cortisol response to exogenous stimuli begins on 4 dpf (Fig. 1A; 29, 30).

**Fig. 1.**
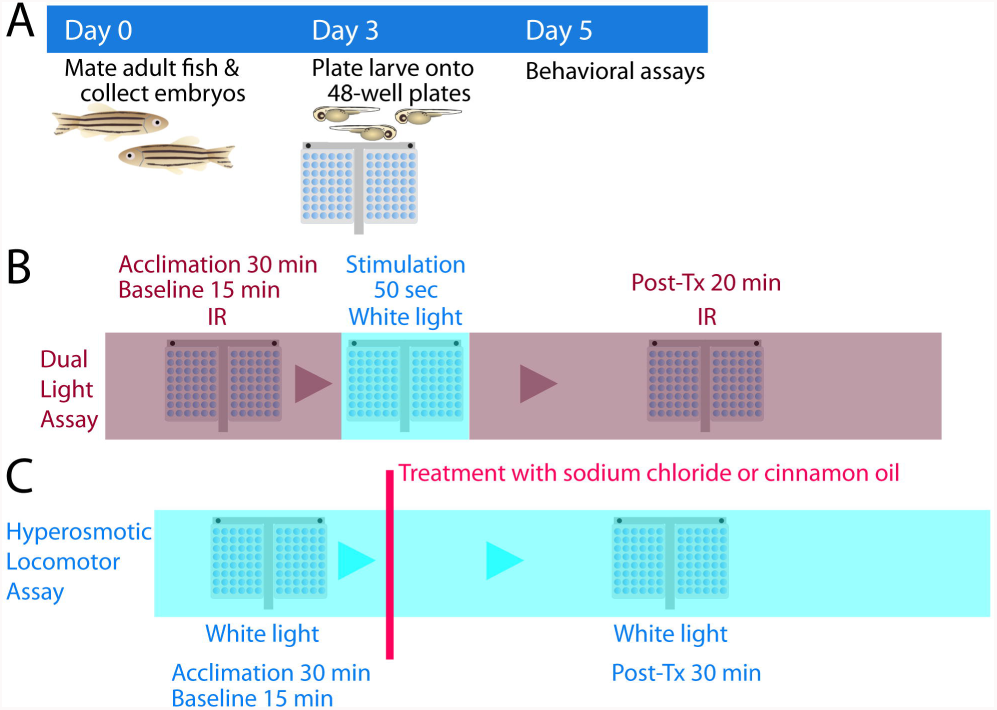
Stress assay paradigms. (*A*) Experimental flow. Embryos are collected from natural spawning on day 0. Dead embryos are cleaned up and fresh embryo media is provided on 0 and 1 dpf. On 3 dpf, morphologically normal larval fish are plated onto 48-well plates. On 5 dpf, stress assays are performed. (*B*) Dual light assays. Larvae are acclimated for 30 min in infrared (IR) light. Baseline locomotor activity is recorded in IR for 15 min, followed by 50 seconds of white light stimulation. Post-treatment locomotor activity is recorded in IR for 20 min. (*C*) Hyperosmotic stress assays. Larvae are acclimated for 30 min in white light. Baseline locomotor activity is recorded for 15 min, followed by addition of NaCl or cinnamon oil (noxious stimulant control). Post-treatment locomotor activity is recorded for 30 min. Initial 10 min were used for statistical analysis for cinnamon oil assays. dpf: days post fertilization.

The dual light assay is developed based on fish visual behavior (Fig. 1*B*). Data from De Marco *et al.*, suggested that non-transgenic control fish that were first dark acclimated and then exposed to blue or yellow light, like transgenics with an optogenetically-induced proopiomelanocortin (*pomc*) gene, resulted in increased locomotion and cortisol release (31). We followed up on these observations with the goal of developing a new assay paradigm for acute stress. The novel dual light assay takes advantage of the fact that zebrafish are essentially blind in an environment illuminated only with infrared (IR) light (32, 33) and exposes fish to white light, which we expected to increase locomotion through activation of the HPI axis as either yellow or blue light did.

The hyperosmotic stress assay is based on fish osmoregulation (Fig. 1*C*). Zebrafish, a freshwater teleost, depend on cortisol synthesis and secretion to excrete extra ions and maintain homeostasis when higher osmotic conditions are encountered (34, 35). We previously reported that larval zebrafish (4 dpf) display increased frequencies of locomotion (number of movement / min) in response to hyperosmotic stress (28). Others have observed that larval zebrafish swim away from an area with an increased osmolarity (36).

A noxious stimulant assay, using cinnamon oil (7.41 µg/mL; ~50 µM), has been used as a control paradigm to show that changes in locomotion are not due to a simple loss of locomotor capacity (Fig. 1*C*). Cinnamon oil is a natural product that is detected by transient receptor potential ion channel ankyrin 1 (*TRPA1*) in mammals that responds to pain inducing noxious stimuli such as the active ingredients in garlic and wasabi (37). Cinnamon oil is a chemical irritant to zebrafish, detected by *trpa1b* expressed in sensory neurons that innervate skin cells, and elicits rapid escape response leading to increased locomotion (38–40).

### Abrupt light change increases locomotor response in wild-type larvae

Because larval zebrafish display clear phototaxic behaviors and wavelength preference (31, 41, 42), we hypothesized that abrupt changes in light illumination are unexpected disruptions that are interpreted as a stressor to induce hypothalamic-pituitary-interrenal (HPI) axis activity and locomotor activity. To test the hypothesis, we challenged wild-type (WT) larvae (5 dpf) with abrupt changes in light conditions (IR-white-IR) with varying lengths of white light (15, 30, 60, 600, or 1800 sec). Significant differences in locomotor response were observed based on duration of (15, 30, 60, 600, or 1800 sec) and exposure to (pre *v*. post-illumination) white light (2-way interaction; *F*_4,1111_ = 15.45, *p* < 0.0005; Fig. 2*A*). Locomotor response significantly increased post-light exposure (main effect; *F*_1,1111_ = 796.41, *p* < 0.0005) and significantly differed based on the durations of white light (main effect; *F*_4,1111_ = 42.33, *p* < 0.0005). Shorter periods of white light stimulation (15, 30, and 60 sec) produced a locomotor profile consisting of two distinct peaks, the first peak occurring before 5 min and the second peak occurring between 10 and 15 min. Speculating that the two peaks may result from distinct signaling cascades, like the sympathetic nervous system activation followed by the HPA axis afterwards, we chose to use 50-seconds of white light illumination as a stimulus for the rest of the experiments so that we could interpret distinct signaling pathways in future investigations.

**Fig. 2.**
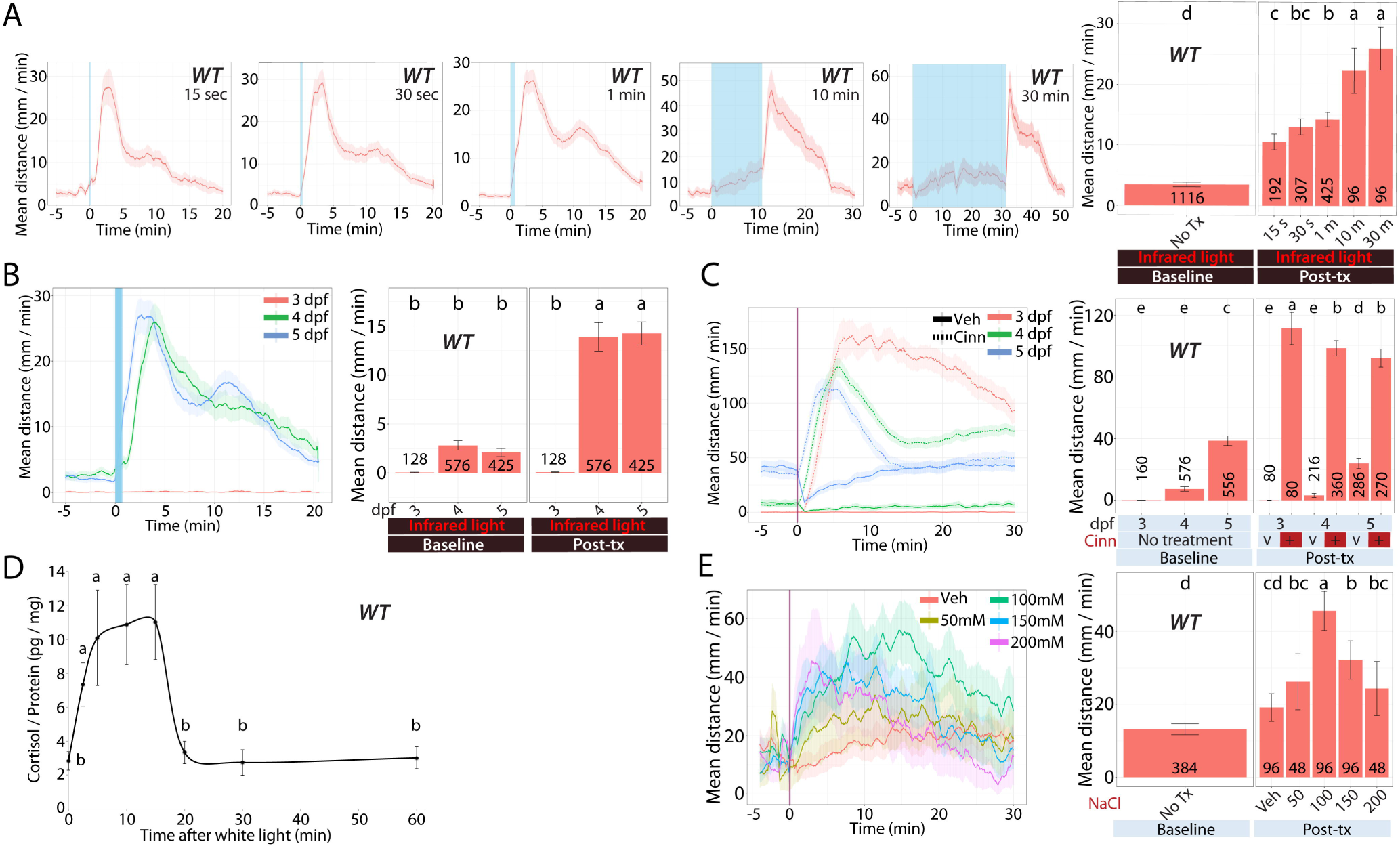
Wild-type larval zebrafish response to acute stressors. We examined the acute response of larvae derived from natural crosses of WT zebrafish in several assays (*A*) Dual light assays with varied lengths of white light illumination (shown as blue bar with times 15, 30, 60, 600, or 1800 sec). All larvae were 5 dpf. (*B*) Dual light assays with larvae at different developmental stages (3, 4, or 5 dpf) after a 50-sec white light illumination. (*C*) Cinnamon oil control assays on (3, 4, or 5 dpf) larvae. (*D*) Whole body cortisol levels in larvae (5 dpf) after a 50-sec white light illumination (*E*) NaCl assays with varying NaCl concentrations on 5 dpf larvae (5 dpf). Line graphs in A, B, C, and E show the rolling mean of the larval distance moved. The locomotor activity at each second is the mean distance fish moved during the preceding 60 s [mean ±95%CI (shading)]. Bar graphs in A, B, C, and E show the mean distance larvae moved over the time course (mm/min; mean ±95%CI) [baseline: 5 min; post-treatment: 20 min (light assays), 30 min (NaCl assays), or 10 min (cinnamon oil control assays)]. The line graph in D shows pg cortisol per mg of total protein over time following a 50-sec white light exposure at time 0. In bar graphs, different letters indicate a significant difference between groups (Tukey’s honest significant difference test, *P* < 0.05). The number of individual larvae measured is shown at the base of each bar graph. WT: wild-type

In the ensuing experiments with 50-seconds of white light illumination, we explored the effect of developmental stages. WT larvae on 4 and 5 dpf, but not on 3 dpf, showed a significantly increased locomotor activity during the post-exposure period in infrared light (Fig. 2*B*). Locomotor response significantly differed by the two-way interaction of the larval age (3, 4, or 5 dpf) and exposure (pre *v*. post-illumination) (*F*_2,1126_ = 38.32, *p* < 0.0005). Locomotion was significantly different based solely on larval age (main effect; *F*_2,1126_ = 52.99, *p* < 0.0005) or on exposure (main effect; *F*_1,1126_ = 227.62, *p* < 0.0005).

When larval zebrafish on 3, 4, or 5 dpf were challenged with cinnamon oil, all age groups showed significantly increased locomotor response post cinnamon oil treatment (Fig. 2*C*). In addition, larvae on 3 dpf showed the most robust locomotor response to cinnamon oil challenge. Larval age resulted in significantly different locomotor response to treatment and exposure [3-way interaction among larval age, treatment (VEH *v*. cinn), and the exposure (pre *v*. post-exposure to the treatment); *F*_2,1286_ = 16.29, *p* < 0.0005]. All 2-way interactions were significant [time and treatment (*F*_1,1286_ = 1037.71, *p* < 0.0005), time and larval age (*F*_2,1286_ = 61.85, *p* < 0.0005), or larval age and treatment (*F*_2,1286_ = 20.21, *p* < 0.0005)].

### Abrupt light change increases whole-body cortisol levels in WT larvae

The stress response (SR) system develops early during embryonic development in zebrafish enabling the HPI axis to respond to exogenous stimuli with cortisol secretion beginning on 4 dpf (29). After observing that abrupt light change increases locomotion in larval zebrafish from 4 dpf on, we tested whether such an increase is correlated with increased whole-body cortisol levels. After a 50-sec white light illumination, larval fish (5 dpf) samples are collected at 2.5, 5, 10, 15, 20, 30, and 60 min after the white light illumination was initiated. The whole-body cortisol levels were significantly increased at 2.5, 5, 10, and 15 min time points compared to that of the control group (0 min) (one-way ANOVA; *F*_7,28_ = 22.76, *p* < 0.0005; Fig. 2*D*). The increased whole-body cortisol levels returned to a value equivalent to that of control samples by 20-min. The significant increase in whole-body cortisol levels from 2.5 to 15 min coincides with the increased locomotor activity that begins within 2 min of light stimulation and lasts for about 15 min (Fig. 2 *A* and *B*).

### Salinity change increases locomotor response in WT larvae

We quantified the total distance travelled by larval zebrafish (5 dpf) after acute sodium chloride application. Our hyperosmotic stress assay confirms that larvae treated with sodium chloride display significantly increased locomotor activities and that the increase in locomotion in response to sodium chloride is concentration-dependent (Fig. 2*E*). Locomotor response significantly differed based on the salt concentration (VEH, 50, 100, 150, or 200 mM) and exposure (pre *v*. post-exposure to the treatment) (2-way interaction; *F*_4,379_ = 12.76, *p* < 0.0005). Locomotor response was significantly different based on the salt concentration (main effect; *F*_4,379_ = 11.28, *p* < 0.0005) or on the exposure (main effect; *F*_1,379_ = 135.55, *p* < 0.0005). We previously showed that acute sodium chloride (100 mM) application increases whole-body cortisol levels (43). Importantly, the 30-min temporal profiles of increased locomotion in this report and increased cortisol levels in our previous report are congruent, showing the peak response at about 15 min after sodium chloride application [(Fig. 2*E*; 43 (Fig. 5*A*)]. Both the behavioral response and the cortisol response occur slightly slower in the hyperosmotic assay relative to the dual light assay.

### Rapid locomotor response to acute stressors is decreased in *mc2r^-/-^* mutant larvae

The association between increased locomotion and increased whole-body cortisol levels suggests a link between cortisol signaling and the observed rapid locomotor response. We tested this hypothesis by blocking systemic glucocorticoid synthesis by knocking out *mc2r* (adrenocorticotropic hormone (ACTH) receptor). *mc2r* is a member of melanocortin receptor family, expressed in interrenal cells at high levels and detectable in adipose tissue (17, 44). *mc2r* gene products specifically bind to ACTH while other melanocortin receptor members (e.g. *mc1r, mc3r*–*mc5r*) bind to melanocyte stimulating hormone as well as ACTH (45–47). *mc2r* is the key molecule that initiates corticosteroid synthesis by increasing cyclic adenosine monophosphate (cAMP) levels, which in turn increases free cholesterol levels available for mitochondrial transport where corticosteroid synthesis occurs (48).

We produced three *mc2r* exon 2 mutant zebrafish alleles (*mc2r^mn57^*, *mc2r^mn58^*, or *mc2r^mn59^*) (Table S2). All three alleles are frame-shift mutations, two 4- and one 5-base pair deletion, which are expected to result in truncated proteins. When 5-dpf larval zebrafish, obtained by in-crossing heterozygous parents, were subject to abrupt light changes (15 min in IR, 50-sec in white light, and 20 min in IR), *mc2r* homozygous mutant siblings showed a significantly decreased locomotor response compared to that of WT siblings (Fig. 3*A*). Locomotor response was significantly different based on the genotype (WT, het, or hom) and exposure (2-way interaction; *F*_2,286_ = 18.82, *p* < 0.0005). Locomotor response was significantly different based on the genotype (main effect; *F*_2,286_ = 4.88, *p* = 0.008) or on the exposure (main effect; *F*_1,286_ = 166.07, *p* < 0.0005). WT and het siblings showed the peak response within a 5-min window and a smaller peak between 10 and 15 min window after a 50-sec of white light illumination (Fig. 3*B*). Both observed peaks are impacted by loss of *mc2r*.

**Fig. 3.**
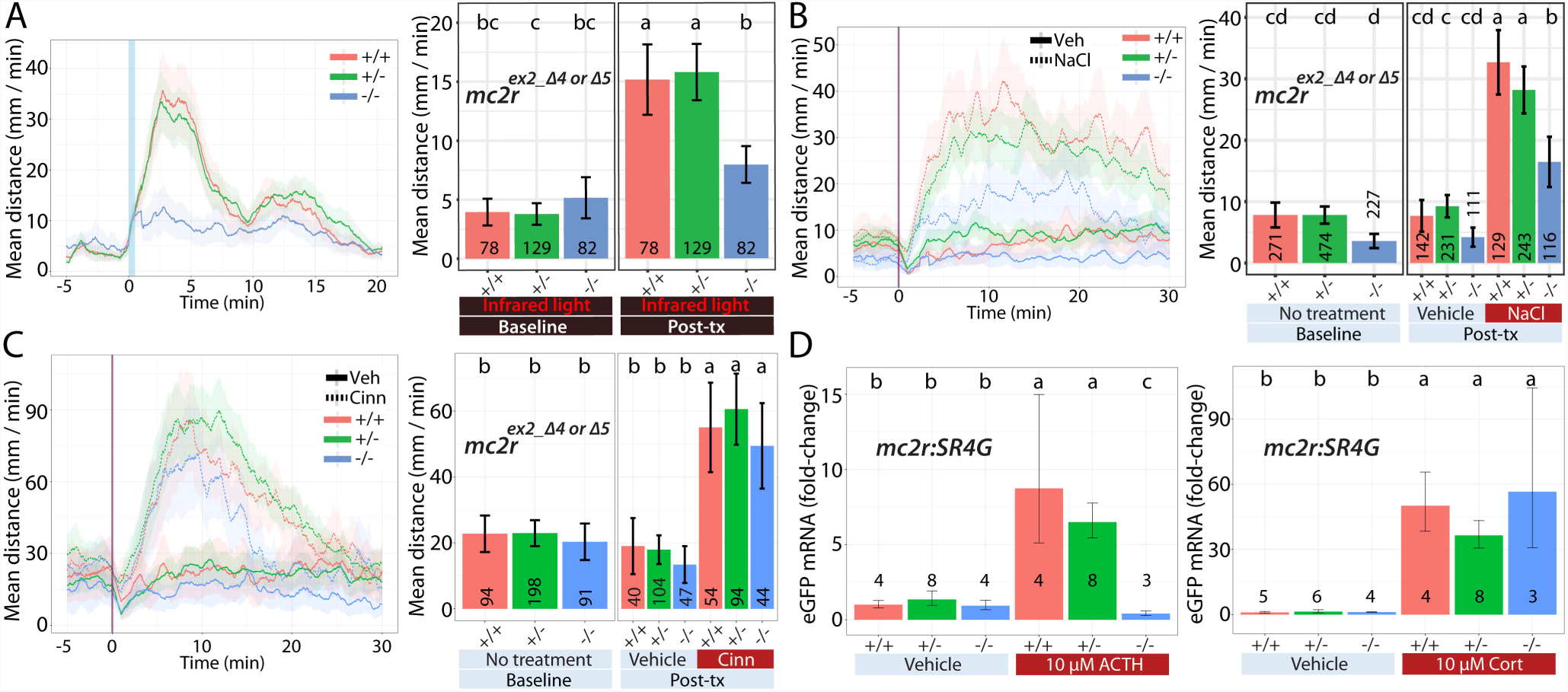
*mc2r* larvae: Locomotor response to acute stressors. We examined the acute response of larvae derived from natural crosses of *mc2r^+/-^* fish [WT (+/+), heterozygous (+/-), or homozygous (-/-)] in several assays. (*A*) Dual light assays, (*B*) NaCl assays, (*C*) Cinnamon oil control assays, and (*D*) Treatment with ACTH or cortisol. The larvae in (*D*) carried a single copy of the SR4G transgene. Line graphs in A, B, and C show the rolling mean of the larval distance moved (5 dpf). The locomotor activity at each second is the mean distance fish moved during the preceding 60 s [mean ±95%CI (shading)]. Bar graphs in A, B, and C show the mean larval distance moved over the time course (mm/min; mean±95%CI) [baseline: 5 min; post-treatment: 20 min (light assays), 30 min (NaCl assays), or 10 min (cinnamon oil control assays)]. Bar graphs in (*D*) show relative EGFP transcript levels compared to WT treated with vehicle. In bar graphs, different letters indicate a significant difference between groups (Tukey’s honest significant difference test, *P* < 0.05). The number of individual larvae measured is shown at the base of each bar graph. SR4G: stress responsive 4-hour half-life GFP

The necessity of glucocorticoid synthesis for rapid locomotor response was examined in the context of sudden salinity changes. When we challenged *mc2r* fish (WT, het, or hom) with 100 mM sodium chloride, *mc2r* homozygous mutant siblings showed a severely decreased, near significant (*p* value between 0.05 and 0.1), locomotor response compared to that of WT siblings (Fig. 3*B*). Locomotor response was near significantly different based on the genotype, treatment (VEH *v*. NaCl) and exposure (3-way interaction; *F*_2,966_ = 2.93, *p* = 0.054). There were significant two-way interactions in all combinations of two factors [genotype and exposure (*F*_2,966_ = 3.89, *p* = 0.021), treatment and exposure (*F*_1,966_ = 129.47, *p* < 0.0005), or genotype and treatment (*F*_2,966_ = 4.13, *p* = 0.016)]. These findings further demonstrate that glucocorticoid synthesis and HPA axis output are critical factors for locomotor response to abrupt salinity changes.

To assure that the decrease in locomotion observed in *mc2r* homozygous mutants is a function of the acute stress response rather than skeletomuscular defects, we performed cinnamon oil assays. Upon cinnamon oil challenge, *mc2r* homozygous mutants showed a comparable locomotor response to that of WT siblings for the first 10-min window, demonstrating preservation of their locomotor capacity (Fig. 3*C*). Locomotor response did not significantly differ based on the genotype, treatment (VEH *v*. cinn), and exposure (3-way interaction; *F*_2,377_ = 0.20, *p* = 0.821). Treatment effect on locomotor response was significant pre or post-exposure (2-way interaction; *F*_1,377_ = 41.53, *p* < 0.0005). Locomotor response was not significantly modified by other two way interactions [genotype and exposure (*F*_2,377_ = 0.40, *p* = 0.668) or genotype and treatment (*F*_2,377_ = 0.72, *p* = 0.489)]. This shows that there was no difference in locomotor response across genotypes in response to the treatment (VEH *v*. cinn) pre or post-exposure during the initial 10-min window.

Since *mc2r* is the key receptor for on-demand cortisol synthesis in response to ACTH signaling, we hypothesized that HPI axis activation in *mc2r* homozygous mutant fish is blocked due to compromised cortisol synthesis. To test the hypothesis, we generated an *mc2r^+/-^*:SR4G^+/-^ transgenic zebrafish strain (*mc2r* heterozygous:SR4G transgene carrier) by crossing *mc2r^mn57^* heterozygous fish with the stress reporter SR4G zebrafish strain (stress responsive 4-hour half-life GFP), which we have previously characterized and reported (43). Briefly, expression of short half-life enhanced green fluorescent protein (EGFP) is driven by transcriptionally active GRs binding to synthetic glucocorticoid response elements (GREs). As a result, when stressed, SR4G fish produce EGFP and transcript levels can be used as a surrogate for HPI axis activation and activated GRs. We treated 5-dpf larvae, obtained by in-crossing *mc2r^mn57^*:SR4G^+/-^ fish, with ACTH (10 µM) or cortisol (10 µM) and quantified the levels of EGFP transcripts with quantitative reverse transcription PCR (qRT-PCR). Following ACTH treatment, EGFP transcript levels were significantly different based on the genotype (WT, het, or hom) and treatment (VEH *v*. ACTH) (2-way ANOVA; *F*_2,25_ = 56.02, *p* < 0.0005; Fig. 3*D*). In addition, ACTH-treated WT siblings had a significantly higher level of EGFP transcripts compared to ACTH-treated homozygotes [*t* test; genotype (WT):treatment (ACTH) *v*. genotype (hom):treatment (ACTH); *t* = 16.34, *p* < 0.0005]. On the contrary, in the cortisol experiment, whereas there was a significant two-way interaction of genotype and treatment (VEH *v*. cortisol) (2-way ANOVA; *F*_2,24_ = 3.87, *p* = 0.035), there was no significant difference between WT and homozygous siblings in two group comparison [*t* test; genotype (WT):treatment (CORT) *v*. genotype (hom):treatment (CORT); *t* = - 0.73, *p* = 0.51].

### Rapid locomotor response to abrupt light change is decreased in *nr3c1* mutant zebrafish

After confirming that increased locomotor response to acute stressors (light or sodium chloride) is dependent, significantly or near significantly, on rapid cortisol synthesis via *mc2r* and subsequent GC signaling, we asked whether or not GC signal transmission requires the canonical glucocorticoid receptor (GR; *nr3c1*). We generated canonical glucocorticoid receptor (*nr3c1*; corticosteroid receptor type II) mutant zebrafish strains by targeting exon 2 (*nr3c1^mn61^* and *nr3c1^mn62^*) or exon 5 (*nr3c1^mn63^*, *nr3c1^mn64^*, and *nr3c1^mn65^*) of the *nr3c1* gene with TALENs (Table S2). The frame-shift mutations are expected to result in truncated proteins. When 5-dpf larvae, obtained by in-crossing heterozygous parents, were subject to abrupt light changes, *nr3c1* homozygous mutant siblings showed a significantly decreased locomotor response compared to that of WT siblings (Fig. 4 *A* and *B*). Locomotor response differed significantly based on the genotype and exposure (*nr3c1^mn61^* and *nr3c1^mn62^*; exon 2; 2-way interaction; *F*_2,449_ = 21.93, *p* < 0.0005). Locomotor response was significantly different based on the genotype (main effect; *F*_2,449_ = 12.06, *p* < 0.0005) or exposure (main effect; *F*_1,449_ = 779.59, *p* < 0.0005).

**Fig. 4.**
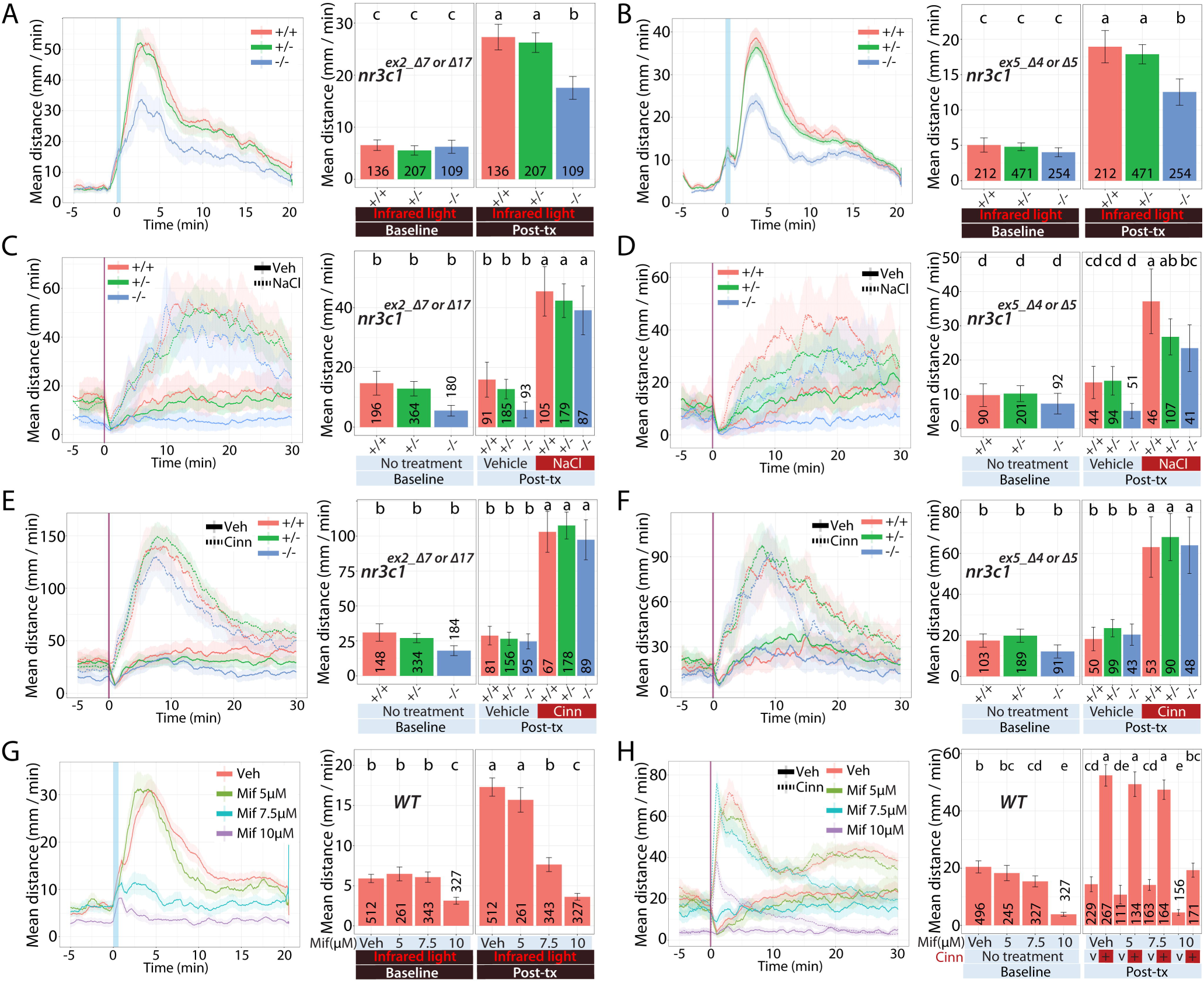
*nr3c1* larvae: Locomotor response to acute stressors. We examined the acute response of larvae derived from natural crosses of *nr3c1^+/-^* fish [WT (+/+), heterozygous (+/-), or homozygous (-/-)] in several assays. Dual light assays for (*A*) *nr3c1^ex2^* and (*B*) *nr3c1^ex5^*. NaCl assays for (*C*) *nr3c1^ex2^* and (D) *nr3c1^ex5^*. Cinnamon oil control assays for (E) *nr3c1^ex2^* and (F) *nr3c1^ex5^*. (*G*) Dual light assays and (*H*) cinnamon oil assays following incubation with mifepristone, a GR antagonist. Line graphs in (*A*)-(*H*) show the rolling mean of the larval distance moved (5 dpf). The locomotor activity at each second is the mean distance fish moved during the preceding 60 s [mean ±95%CI (shading)]. Bar graphs in (*A*)-(*H*) show the mean distance larvae moved over the time course (mm/min; mean±95%CI) [baseline: 5 min; post-treatment: 20 min (light assays), 30 min (NaCl assays), or 10 min (cinnamon oil control assays)]. In bar graphs, different letters indicate a significant difference between groups (Tukey’s honest significant difference test, *P* < 0.05). The number of individual larvae measured is shown at the base of each bar graph. GR: glucocorticoid receptor

Locomotor response (*nr3c1^mn63^*, *nr3c1^mn63^*, and *nr3c1^mn63^*; exon 5) was significantly different based on the genotype and exposure (2-way interaction; *F*_2,934_ = 11.46, *p* < 0.0005). Locomotor response significantly differed based on the genotype (main effect; *F*_2,934_ = 11.22, *p* < 0.0005) or exposure (main effect; *F*_1,934_ = 606.71, *p* < 0.0005). The increase in locomotion in WT and het siblings occurs within 5 min after a 50-sec exposure to white light (Fig. 4 *A* and *B*). This observation supports that locomotor response to abrupt light change requires rapid GC-GR signaling via the canonical GR (*nr3c1*).

While the canonical GR (*nr3c1*) is required to respond to sudden light changes, it does not as dramatically impact changes measured following sudden salinity changes. When 5-dpf larvae— WT, het, and hom siblings at *nr3c1* exon 2 or 5—were challenged with 100 mM sodium chloride, *nr3c1^ex5^* homozygous mutants showed a near significant decrease in locomotor response compared to that of WT siblings, whereas *nr3c1^ex2^* homozygous siblings showed a moderate, but non-significant, reduction in locomotor response (Fig. 4 *C* and *D*).

Locomotor response (*nr3c1^ex2^*) did not differ based on the genotype, treatment (VEH *v*. NaCl), and exposure (3-way interaction; *F*_2,734_ = 0.1, *p* = 0.907). Locomotor response significantly differed based on treatment and exposure (2-way interaction; *F*_1,734_ = 185.94, *p* < 0.0005). Other factors did not significantly alter locomotor response [2-way interactions; genotype and exposure (*F*_2,734_ = 0.34, *p* = 0.713) or genotype and treatment (*F*_2,734_ = 0.23, *p* = 0.80)]. Locomotor response (*nr3c1^ex5^*) differences were near significant based on the genotype, treatment, and exposure (3-way interaction; *F*_2,377_ = 2.45, *p* = 0.088). Locomotor response significantly differed based on the treatment and exposure (2-way interaction; *F*_1,377_ = 62.14, *p* < 0.0005). Other factors did not significantly modify locomotor response [2-way interactions; genotype and exposure (near significant) (*F*_2,377_ = 2.82, *p* = 0.061) or genotype and treatment (*F*_2,377_ = 0.67, *p* = 0.51)]. Locomotor response in *nr3c1* exon 5 differed with a greater magnitude based on the genotype, treatment, and exposure compared that in *nr3c1* exon 2.

Upon cinnamon oil challenge, *nr3c1* homozygous mutants (both exon 2 and 5) showed a comparable locomotor response to that of WT siblings within the initial 10-min window, demonstrating their locomotor capacity was preserved (Fig. 4 *E* and *F*). Locomotor response (*nr3c1^ex2^*) to cinnamon oil challenges did not differ based on the genotype, treatment, and exposure (3-way interaction; *F*_2,660_ = 2.06, *p* = 0.128). Locomotor response significantly differ based on the treatment and exposure (*F*_1,660_ = 296.57, *p* < 0.0005). Other factors did not modify locomotor response [2-way interactions; genotype and exposure (*F*_2,660_ = 0.94, *p* = 0.392) or genotype and treatment (*F*_2,660_ = 0.07, *p* = 0.932)]. Similarly, locomotor response (*nr3c1^ex5^*) did not differ based on the genotype, treatment, and exposure (3-way interaction; *F*_2,377_ = 0.14, *p* = 0.869). Locomotor response significantly differed based on the treatment and exposure (2-way interaction; *F*_1,377_ = 87.91, *p* < 0.0005). Other factors did not modify locomotor response [2-way interactions; genotype and exposure (*F*_2,377_ = 0.61, *p* = 0.542) or genotype and treatment (*F*_2,377_ = 0.15, *p* = 0.861)]. The cinnamon oil assays for the alleles on *nr3c1^ex2^* and *nr3c1^ex5^* demonstrate that there was no difference in locomotion due to genotype in response to the treatment at pre or post-exposure, and that the locomotor capacity of these mutants was not impaired.

We hypothesized that, if decreased locomotor response to abrupt light change in *nr3c1* homozygotes is dependent on *nr3c1*, a canonical GR antagonist, mifepristone (RU38486), would block the stressor stimulated locomotion in WT fish replicating our findings with *nr3c1* loss-of-function alleles. Treating WT fish (5 dpf) with varying doses of mifepristone (5, 7.5, or 10 µM) resulted in significantly decreased locomotor responses at 7.5 µM, a dose that still maintained a rapid responsiveness to cinnamon oil (Fig. 4 *G* and *H*). At 5 µM, locomotor response was comparable to VEH-treated fish whereas 10 µM mifepristone significantly decreased locomotion in both abrupt light change and cinnamon oil challenge.

In light assays, locomotor response significantly differed based on the mifepristone dose (VEH, 5, 7.5, or 10 µM) and exposure (2-way interaction; *F*_3,1439_ = 132.50, *p* < 0.0005). In addition, locomotion significantly differed based on the mifepristone dose (main effect; *F*_3,1439_ = 107.82, *p* < 0.0005) or exposure (main effect; *F*_1,1439_ = 508.30, *p* < 0.0005). Similarly, in cinnamon oil assays, locomotor response significantly differed based on the mifepristone dose, treatment (VEH *v*. cinn), and exposure (3-way interaction; *F*_3,1387_ = 16.30, *p* < 0.0005). Locomotion significantly differed based on all combinations of two-way interactions [treatment and exposure (*F*_1,1387_ = 561.21, *p* < 0.0005), mifepristone dose and exposure (*F*_3,1387_ = 6.38, *p* < 0.0005), or mifepristone dose and treatment (*F*_3,1387_ = 11.6, *p* < 0.0005)].

These outcomes show that 7.5 µM mifepristone, a dose that does not impact locomotor response to cinnamon oil challenges as compared to VEH-treated fish [*t* test; mif dose (VEH):treatment (cinn):exposure (post) *v*. mif dose (7.5 µM):treatment (cinn):exposure (post); *t* = -0.12, *p* = 0.91], inhibits locomotor response to light illumination changes [*t* test; mif dose (VEH):exposure (post) *v*. mif dose (7.5 µM):exposure (post), *t* = -13.24, *p* < 0.0005]. These results further reinforce that the canonical GR (*nr3c1*) is required for locomotor response to abrupt light change.

### Rapid locomotor response to acute stressors is not decreased in *nr3c2* mutants

After confirming the involvement of the canonical GR in rapid locomotor response to abrupt light change, we tested whether or not mineralocorticoid receptor (MR; corticosteroid receptor type I) is required for the response. MR encoded by *nr3c2* is highly expressed in the limbic areas of the brain, has a 10-fold higher affinity for glucocorticoids than the canonical GR, and may be plasma membrane associated (49, 50).

Using TALENs, we generated MR mutant zebrafish strains in exon 2 (*nr3c2^mn66^* and *nr3c2^mn67^*) of the *nr3c2* gene (Table S2). When 5-dpf larvae were subject to abrupt light changes, *nr3c2* homozygous mutant siblings showed a comparable locomotor response to that of WT siblings (Fig. 5*A*). Locomotor response did not differ based on the genotype and exposure (2-way interaction; *F*_2,431_ = 0.22, *p* = 0.805). Locomotor response significantly increased across all genotype post exposure to white light (main effect; *F*_1,431_ = 500.53, *p* < 0.0005). Locomotor response did not differ based on the genotype (main effect; *F*_2,431_ = 0.65, *p* = 0.524).

**Fig. 5.**
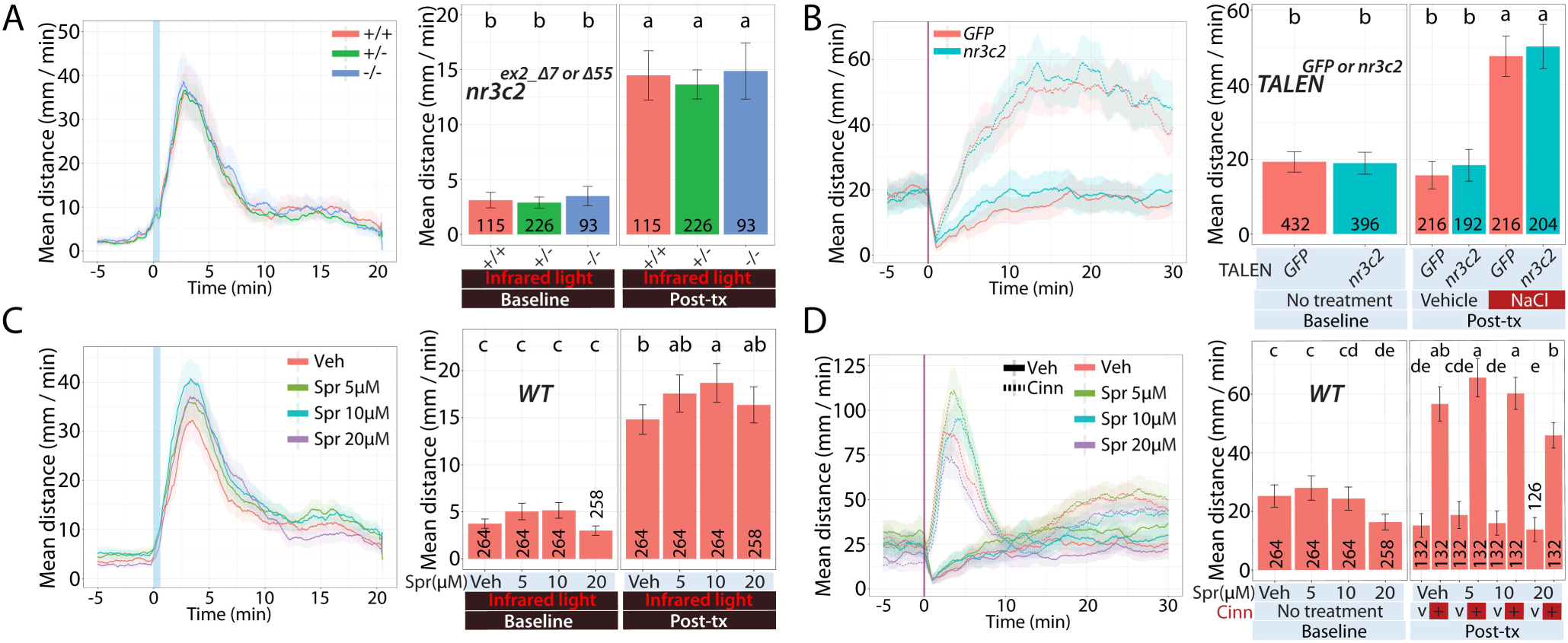
*nr3c2* larvae: Locomotor response to acute stressors. We examined the acute response of larvae derived from natural crosses of *nr3c2^+/-^* fish [WT (+/+), heterozygous (+/-), or homozygous (-/-)] or injected with high efficiency, bi-allelic TALENs targeting GFP sequences or *nr3c2* exon 2 in several assays. (*A*) Dual light assays for *nr3c2* larvae. (*B*) NaCl assays for TALEN-injected WT larvae. (*C*) Dual light assays or (D) cinnamon oil assays following incubation with spironolactone, a MR antagonist. Line graphs in (*A*)-(*D*) show the rolling mean of the larval distance moved (5 dpf). The locomotor activity at each second is the mean distance fish moved during the preceding 60 s [mean ±95%CI (shading)]. Bar graphs in (*A*)-(*D*) show the mean distance larvae moved over the time course (mm/min; mean±95%CI) [baseline: 5 min; post-treatment: 20 min (light assays) or 30 min (NaCl assays)]. In bar graphs, different letters indicate a significant difference between groups (Tukey’s honest significant difference test, *P* < 0.05). The number of individual larvae measured is shown at the base of each bar graph. MR: mineralocorticoid receptor, TALEN: transcription activator like effector nuclease

Consistent with the results of germline mutant fish in dual-light assays, when 5-dpf WT larvae injected with high-efficiency, biallelic TALENs targeting *nr3c2* exon 2 were challenged with 100 mM sodium chloride, *nr3c2^ex2_TALEN_inj^* fish showed a comparable locomotor response to that of WT siblings sham-injected with GFP sequence targeting TALENs (Fig. 5*B*). Locomotor response did not differ based on the injection reagent (VEH *v*. *nr3c2*-targeting TALEN), treatment (VEH *v*. NaCl), and exposure (3-way interaction; *F*_1,824_ = 0.09, *p* = 0.77). Locomotor response significantly differed based on the treatment and exposure (2-way interaction; *F*_1,824_ = 163.72, *p* < 0.0005). Other factors did not alter locomotor response [2-way interactions; injection reagent and exposure (*F*_1,824_ = 1.44, *p* = 0.231) or injection reagent and treatment (*F*_1,824_ = 0.027, *p* = 0.87)].

We hypothesized that, if locomotor response to abrupt light change does not require *nr3c2*, an MR antagonist, spironolactone, would not block the locomotion in WT fish replicating the finding. Treating WT larval fish (5 dpf) with varying doses of spironolactone (5, 10, or 20 µM) resulted in a slight, but significantly increased locomotor response at 10 µM that did not significantly change responsiveness to cinnamon oil (Fig. 5*C*).

In light assays, locomotor response did not differ based on the drug dose (VEH, 5, 10, or 20 µM) and exposure (2-way interaction; *F*_3,1046_ = 1.57, *p* = 0.196). However, locomotor response differed based on the drug dose (main effect; *F*_3,1046_ = 5.05, *p* = 0.002) or exposure (main effect; *F*_1,1046_ = 788.06, *p* < 0.0005). In cinnamon oil assays, locomotor response significantly differed based on the drug dose, treatment, and exposure (3-way interaction; *F*_3,1042_ = 2.94, *p* = 0.032).

Locomotor response significantly differed based on the treatment and exposure (2-way interaction; *F*_1,1042_ = 471.83, *p* < 0.0005). Other two-way interactions did not alter locomotor response [drug dose and exposure (*F*_3,1042_ = 0.64, *p* = 0.587) or drug dose and treatment (*F*_3,1042_ = 1.46, *p* = 0.224)].

At 5 through 20 µM of spironolactone applications, locomotor response to light illumination changes was comparable or slightly increased compared to that of VEH-treated fish. Locomotor response to cinnamon oil challenges produced similar outcomes except that, at 20 µM, the locomotor response was significantly decreased (Fig. 5 *C* and *D*). This observation supports that *nr3c2* is not necessary for locomotor response to abrupt light change at the doses that do not produce more generalized effects on locomotion (shown by response to cinnamon oil challenge comparable to VEH-treated fish).

### Anxiolytic and anxiogenic drugs modulate locomotor response to abrupt light change

To test the translational potential of our acute stress assay suite for screening pharmacological compounds, we tested the predictive validity of the dual light assay paradigm. Predictive validity can be assessed by treating animals with therapeutics prescribed for humans and monitoring whether the ensuing outcomes in the animals fit with the reported effects of the compounds in humans (51).

Paroxetine is a widely prescribed antidepressant (selective serotonin reuptake inhibitor (SSRI)). Paroxetine and antidepressants in general have been shown to normalize HPA axis activity in humans and rodent models in prolonged treatment contexts (52, 53). Although the effect of antidepressant in normalizing HPA axis output is focused on prolonged treatments (e.g. weeks), there is still potential for acute normalizing effects of antidepressants since pleiotropic effects of antidepressants have not been fully understood. We hypothesized that, if increased locomotor response to abrupt light challenge is dependent on HPA axis activation, paroxetine application on WT fish would attenuate the locomotor response because of presumed acute effects of antidepressants in normalizing HPA axis activity.

For light assays, WT larval fish were pre-treated overnight between 3 and 4 dpf with varying doses of paroxetine (0.625 or 1.25 µM) and the drug is removed on 4 dpf. This treatment resulted in significantly decreased locomotor responses to an acute light stressor on 5 dpf (Fig. 6*A*). Locomotor response significantly differed based on the drug dose (VEH, 0.625, or 1.25 µM) and exposure (2-way interaction; *F*_2,1027_ = 19.57, *p* < 0.0005). In cinnamon oil assays (Fig. 6*B*), locomotor response significantly differed based on the drug dose, treatment, and exposure (3-way interaction; *F*_2,1024_ = 6.09, *p* < 0.002). Locomotor response significantly differed based on some combinations of two-way interactions [treatment and exposure (*F*_1,1024_ = 694.43, *p* < 0.0005) or drug dose and exposure (*F*_2,1024_ = 6.48, *p* = 0.002)], There was no significant interaction between drug dose and treatment (*F*_2,1024_ = 2.53, *p* = 0.08). In particular, 0.625 µM paroxetine did not attenuate a rapid locomotor response to cinnamon oil [*t* test; dose (VEH):treatment (cinn):exposure (post) *v*. dose (0.625 µM):treatment (cinn):exposure (post); *t* = 0.18, *p* = 0.86] while decreasing locomotor response in light assays [*t* test; dose (VEH):exposure (post-illumination) *v*. dose (0.625 µM):exposure (post); *t* = 6.24, *p* < 0.0005]. This observation supports that increased locomotor response to abrupt light change is dependent on HPI axis activation and can be attenuated via application of anxiolytic drugs.

**Fig. 6.**
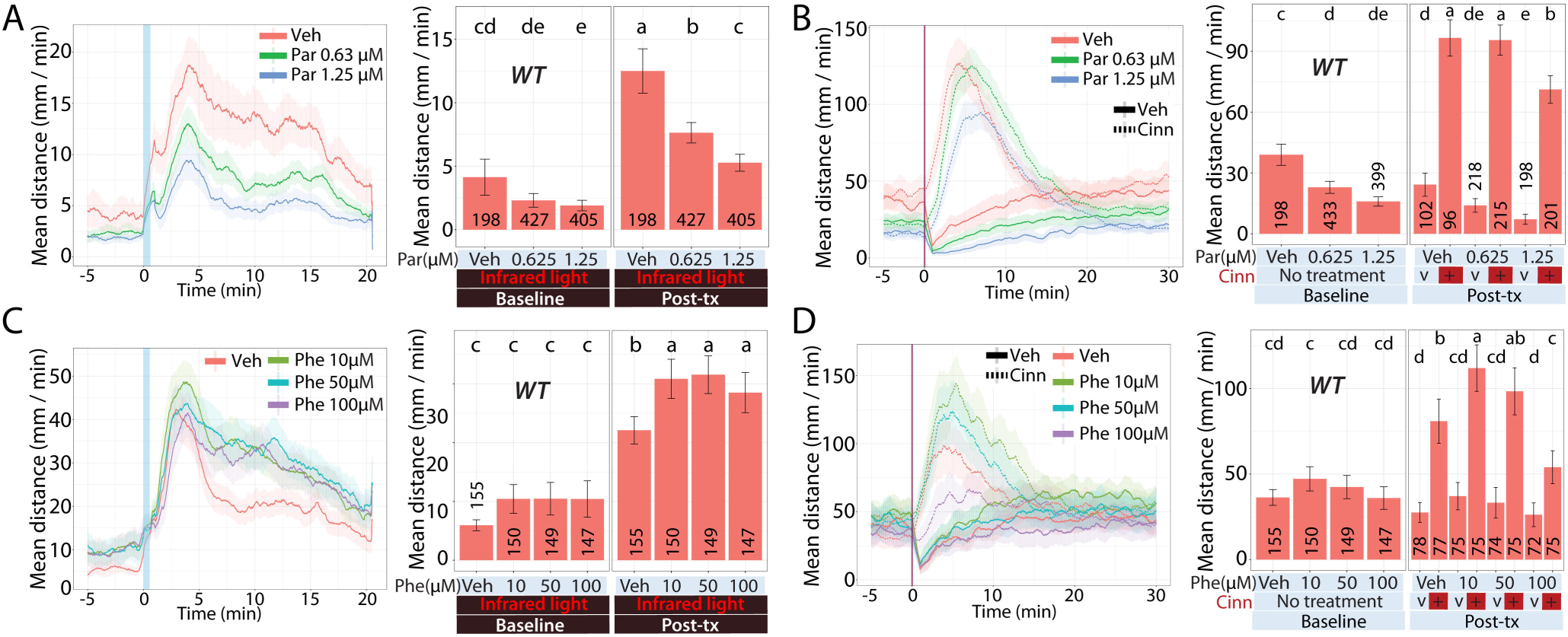
WT larvae pre-incubated with drugs: Locomotor response to acute stressors. We examined the acute response of larvae derived from natural crosses of WT fish, following with an anyxiolytic or anxiogenic drug. (*A*) Dual light assays or (*B*) cinnamon oil assays following incubation with paroxetine, an SSRI. (*C*) Dual light assays or (*D*) cinnamon oil assays following incubation with phenylephrine, an α1 adrenoceptor agonist. Line graphs in (*A*)-(*D*) show the rolling mean of the larval distance moved (5 dpf). The locomotor activity at each second is the mean distance fish moved during the preceding 60 s [mean ±95%CI (shading)]. Bar graphs in (*A*)-(*D*) show the mean distance larvae moved over the time course (mm/min; mean±95%CI) [baseline: 5 min; post-treatment: 20 min (light assays) or 10 min (cinnamon oil control assays)]. In bar graphs, different letters indicate a significant difference between groups (Tukey’s honest significant difference test, *P* < 0.05). The number of individual larvae measured is shown at the base of each bar graph. SSRI: Selective serotonin reuptake inhibitor

We tested the effect of a stimulant, phenylephrine. Phenylephrine is an α1 adrenergic receptor agonist. α1 adrenoceptors are co-expressed with CRH containing neurons in the hypothalamus in rats (54) and have been shown to increase HPA axis output (55, 56). Further, in the nuclei (e.g. nucleus of the solitary tract, locus coeruleus) in the brainstem that provide major noradrenergic projections to the PVN, catecholaminergic lesions or stimulation resulted in attenuated or increased HPA axis output, respectively (57, 58). Catecholamine-mediated stimulation of HPA axis output was α1 adrenergic receptor dependent (59). We hypothesized that, if increased locomotor response to abrupt light challenge is dependent on HPA axis activation, phenylephrine application on WT fish would potentiate the locomotor response because SNS output positively regulates HPA axis activity. Pre-treating WT larval fish on 5 dpf for 2 hours with varying doses of phenylephrine (10, 50, or 100 µM) resulted in significantly increased locomotor responses in light assays at all doses tested (Fig.6*C*). Locomotor response differed based on the drug dose (VEH, 10, 50, or 100 µM phenylephrine) and exposure at near significant levels (2-way interaction; *F*_3,597_ = 2.58, *p* = 0.052). Locomotor response significantly differed based on drug doses (main effect; *F*_3,597_ = 7.27, *p* < 0.0005) or exposure (main effect; *F*_1,597_ = 718.44, *p* < 0.0005).

When challenged with cinnamon oil, only the lowest dose (10 µM) significantly increased locomotor response whereas a comparable response to VEH-treated fish resulted at 50 µM and a significantly decreased response resulted at the highest dose (100 µM; Fig. 6*D*). Locomotor response significantly differed based on the drug dose (VEH, 10, 50, or 100 µM phenylephrine), treatment, and exposure (3-way interaction; *F*_3,593_ = 6.91, *p* < 0.0005). Locomotor response was significantly altered based on all combinations of two factors [treatment and exposure (*F*_1,593_ = 207.56, *p* < 0.0005), drug dose and exposure (*F*_3,593_ = 6.19, *p* < 0.0005), or drug dose and treatment (*F*_3,593_ = 3.22, *p* = 0.022)]. The 50 µM phenylephrine dose significantly increased locomotor response to light changes [*t* test; dose (VEH):exposure (post) *v*. dose (50 µM):exposure (post); *t* = -4.71, *p* < 0.0005] whereas the same dose led to no significant change in locomotor response compared to VEH-treated (0 µM phenylephrine) group [*t* test; dose (VEH):treatment (cinn):exposure (post) *v*. dose (50 µM):treatment (cinn):exposure (post); *t* = - 1.84, *p* = 0.07]. This observation supports that the light assay sensitively reflects changes in HPI output and anxiogenic drugs can potentiate locomotor response to abrupt light change.

## Discussion

We demonstrated that locomotor response to abrupt changes in light illumination or salinity requires HPI axis function, including activation of glucocorticoid synthesis via *mc2r* (ACTH receptor) and canonical glucocorticoid receptor (GR; *nr3c1*) activity. The locomotor response did not require mineralocorticoid receptor (MR; *nr3c2*). Knocking out MR leads to a slightly increased locomotor trend in *nr3c2* exon 2 homozygous siblings. The locomotor response phenotypes of homozygous mutant fish in corticosteroid receptor type II (GR) or I (MR) are replicated in mifepristone (GR antagonist) or spironolactone (MR antagonist) treatment, respectively, leading to a significant decrease or increase in locomotion. Anxiolytic drugs (SSRIs) or anxiogenic compounds (phenylephrine) attenuated or potentiated the locomotor response, respectively, providing supporting evidence that rapid locomotor response to acute stressors are associated with neural pathways processing anxiety-like behaviors and that our assay system has a potential to be used for drug screening for such classes of drugs.

### Diverse variables led to distinct locomotor profiles

Developmental stages are found to be important in locomotor response to acute changes in light illumination as well as in response to noxious stimuli (cinnamon oil). Whereas larval fish at 3 dpf did not show increased locomotion following white light illumination, larvae at 4 or 5 dpf responded to the changes with significantly increased locomotion (Fig. 2*B*). This agrees with the finding that zebrafish do not robustly respond to exogenous stimuli with cortisol secretion until 4 dpf on (96 hpf) even though the HPI axis begins developing after 2 dpf (29, 30). However, when larvae on 3 dpf were challenged with cinnamon oil, they showed the most robust locomotor response compared to larvae on 4 and 5 dpf (Fig. 2*C*). Such differential effects of light illumination or cinnamon oil treatment illustrate the characteristics of the developing HPI axis from 3 to 5 dpf; the more mature HPI activities beginning at 4 dpf enable more sensitive responses (responses to light) and more regulated responses (responses to cinnamon oil).

Different stressor types (light or salinity) produced distinct locomotor profiles. While abrupt light changes elicited a faster increase in locomotion peaking in 5 min after light change, abrupt salinity changes led to increased locomotion that peaked at about 15 min. Different sensory pathways are likely to determine the response profile. Changes in illumination are detected through the visual system while those in osmolarity are via osmoregulatory systems (35, 60). Whereas sensing changes in illumination can be immediate, the time domain required to sense changes in osmolarity outside and inside the body and to initiate physiological and behavioral response such as ion filtration changes and locomotion would be longer.

Varying lengths of white light illumination also produced distinct locomotor profiles. Longer illuminations (10 or 30 min) produced higher peaks compared to shorter illuminations (15, 30 or 60 sec) (Fig. 2*A*). Varying concentrations of sodium chloride produce quantitatively (more or less) and temporally (quicker or slower) different responses. The stimulus that we chose was 100 mM sodium chloride. Seawater is at about 600 mM and freshwater in rivers is at about 1 mM NaCl (actual salinities are calculated based on many different salts) (61, 62). The salinity in a zebrafish body is about 150 mM, which is similar to that in humans (63, 64). Although the concentration (100 mM) in our stimulation is higher than that of freshwater, it is far less than that of seawater and below the physiological salinity in zebrafish body. This would not reverse the normal physiology of zebrafish in which the animal needs to constantly excrete excess water to maintain osmotic and ionic homeostasis (65, 66). Such concentration of 100 mM salt is not highly likely to be overwhelming. In sodium chloride dose curve experiments, 200 mM NaCl challenge induced a quicker response reaching the peak response before 10 min (Fig. 2*E*). NaCl concentrations greater than or equal to 150 mM led to a decreased total locomotor response, compared to those challenged with 100 mM NaCl, which may indicate physiologically detrimental effects at those higher concentrations (150 or 200 mM).

A noxious stimulus (cinnamon oil) produces a distinct locomotor profile to those produced by light illumination and salinity change. While changes in illumination or salinity are not threatening to zebrafish survival, stimulations by cinnamon oil may be perceived threatening as it is sensed through the *trpa1b* channel that mediates noxious and painful stimuli in mammals (*TRPA1*) (37). A unique feature of locomotor responses to cinnamon oil challenges is that the response quickly rose regardless of the genotype and more rapidly subsided in homozygous mutants. For instance, whereas the locomotor response of all homozygous mutants (*mc2r* or *nr3c1^ex5^*) to cinnamon oil challenge was comparable to that of WT and het siblings for the first 10-15 min periods following treatment, the locomotor response in homozygous mutants did not last as long as that of WT or het siblings did (Fig. 3*C* and 4*F*). This quicker tapering of locomotor response in homozygotes may be indicative of a potential involvement of HPA axis activation and glucocorticoid signaling in the continued response to a noxious stimulus. It is also possible that, because our environmental stress assays (light or hyperosmotic assays) utilize mild to moderate levels of stimuli, the variations in response characteristics of individual fish may be revealed. In other words, faced with a stronger stimulus such as cinnamon oil, all fish could be responding uniformly due to the strong saliency of the stimulus whereas milder stressors such as changes in light may induce a wider range of locomotor response between fish. In any case, such robust locomotor response to cinnamon oil during the early window (initial 10 min) demonstrates that *mc2r* or *nr3c1* homozygous mutant siblings have intact locomotor capability.

### Whole-body cortisol levels are congruent with locomotor response profiles

The temporal profiles of locomotion coincided with changes in whole-body cortisol levels. Light changes induced significantly increased cortisol levels from 5 to 15 min after the white light illumination (Fig. 2) while 100 mM sodium chloride application led to peak cortisol levels at 20 min (43). In addition, the rapid decrease in cortisol levels between 15 and 20 min in dual light assays (Fig. 2*C*) implies that there is a mechanism that quickly degrades or excretes cortisol enabling a tight regulation of cortisol availability. After high levels of cortisol are detected, gradual decrease in cortisol levels is reported in zebrafish that takes between 30 min to 2 hrs in larval or adult zebrafish (67, 68). The rate of cortisol degradation *in vivo* is not well established due to the differences in cortisol quantification (e.g. amount per embryo or amount in the trunk of an adult fish). In our studies, the hyperosmotic stress assays with 100 mM sodium chloride showed a degradation rate of about 25 pg body cortisol per min (43) while the dual assays showed approximately 2.5 pg body cortisol per min (Fig. 2*D*). The mechanisms that regulate cortisol bioavailability may be an important area of research in the context of neuropsychiatric disorders.

### mc2r or corticosteroid receptor knock-outs have differential effects

*mc2r* homozygous knock-outs showed significantly decreased locomotor responses in both light and hyperosmotic stress assays (Fig. 3 *A* and *B*). On the other hand, sibling embryos obtained from *nr3c1* heterozygous in-crosses showed more nuanced outcomes. In light assays, significant interactions between the genotype (WT, het, or mom) and exposure (pre *v*. post-exposure) were found in both *nr3c1^ex2^* and *nr3c1^ex5^*. However, in hyperosmotic stress (sodium chloride) assays, there was no significant 3-way interaction among the genotype (WT, het, or hom), treatment type (VEH *v*. NaCl), or exposure (pre *v*. post-exposure) in both *nr3c1^ex2^* and *nr3c1^ex5^*. However, the *p* values were 0.088 for *nr3c1^ex5^* and 0.907 for *nr3c1^ex2^*, showing that there is a stronger interaction effect in *nr3c1^ex5^*.

The difference between *mc2r* and *nr3c1* knockouts may result from the fact that the *mc2r* gene has a simpler regulatory environment than *nr3c1* does, regarding its protein expression. *mc2r* has two exons and only exon 2 is responsible for protein coding. The Mc2r protein is a trans-membrane protein with only one transcript and translational isoform (46, 69, 70). In contrast, *nr3c1* has 9 exons, with which exon 1 and 9 produce three and two alternative transcripts, respectively, and has 7 alternative translational isoforms (71–73). Nr3c1 forms a cytoplasmic complex with chaperone proteins, is involved in various signaling pathways, and functions as a transcription factor (74, 75). The complex environment regulating Nr3c1 function points to the possibility that diverse Nr3c1 isoforms may play differential roles in distinct tasks, tissues, or neural circuits. In this context, our frame-shift mutations in either exon 2 or 5 might not completely eliminate all the gene products and that alternative translational isoforms downstream of a premature stop codon may be still at work in locomotor response to salinity changes but not to changes in light conditions. This line of thought is supported by the fact that 3-way interaction was close to statistical significance in *nr3c1^ex5^* (*p* = 0.088) compared to that in *nr3c1^ex2^* (*p* = 0.907). Alternatively, exon skipping of exon 2 or exon 5 may be occurring to generate alternative *nr3c1* mRNA transcripts and these forms differentially function in response to environment changes. Still, it is also likely that hyperosmotic assays that use salinity changes as the stimulant may involve more organ systems, physiologic processes, and signaling pathways involved in osmotic and ion homeostasis, making the locomotor response to NaCl more diverse compared to dual-light assays.

When GR is genetically knocked out or pharmacologically blocked (mifepristone) between 4 and 5 dpf, the locomotion is significantly attenuated in light assays. Two alternative hypotheses for this observation may be proposed. One is that GR is needed during development to establish the tone of the HPA axis and the other being GR is needed for direct and rapid signaling while the locomotor response is occurring. In rodent models, GR knock-out animals often display depression-like behavioral changes, as well as neuroendocrine abnormalities (76, 77). When GR is knocked out, feedback inhibition by GC is impaired and a constitutively high secretion of CRH, ACTH, and cortisol ensues, which in turn leads to general downregulation of GC receptors, GC resistance, and blunted stress response to external stimuli. In our assay paradigm, the subject fish are 5 dpf, and are naive to stress. Loss of GR in these fish during the development of the HPI axis may result in altered HPA functions and stress response. To tease out the developmental hypothesis with the alternative hypothesis that GR is needed for direct and rapid signaling during the locomotor response, additional experiments are necessary. Conditional zebrafish strains could be useful to knock out GR at a specific time point closer to day 5 to minimize the effect of loss of GR during HPI axis development.

When MR is genetically knocked out or pharmacologically blocked (spironolactone) between 4 and 5 dpf, the locomotion is comparable to that of WT or significantly increased in light assays, respectively. The small but significantly increased locomotor response in spironolactone treatment (Fig. 5*C*) indicates that there is a possibility that a complete deletion of *nr3c2* gene may result in significantly increased locomotor response in homozygous mutants. The current MR homozygous mutants (*nr3c2*^-/-^) are generated from frameshift mutations (7 or 55 del) in exon 2. There may be some residual gene function remaining from alternative transcripts or translational isoforms, like other nuclear hormone receptors including thyroid hormone receptors (78) and *nr3c1* (73).

### Future investigations on rapid GC signaling and stress response

Nongenomic GC signaling drives rapid behavioral changes via the central nervous system within minutes of stimulation, which range from reproductive behavior to response to novel environments. These behavioral changes are observed in a variety of vertebrate species such as amphibians (79, 80), birds (81, 82), and rodents (83, 84) and exemplify the conserved nature of rapid action of GCs and their cognate receptors in animal behaviors. In amphibians (rough-skinned newts), GC signaling that rapidly reduces male reproductive behavior is mediated via a population of medullary neurons in the brainstem (85). In many species, neural tissues and pathways driving those adaptive behavioral changes remain to be identified. We investigated the essential role of rapid GC signaling in the context of locomotor responses to acute changes in the environment. Our findings show that rapid locomotor response is a stress response dependent on HPI axis activation and the canonical glucocorticoid receptor. It is important and now feasible to investigate neural circuits that rapidly modulate the neuroendocrine arm (HPI axis) of stress response and lead to behavioral adaptations in living animals.

At the cellular level, nongenomic GC-CR signaling rapidly alters various aspects of neuronal physiology and functions. GC signaling swiftly (seconds–minutes) modulates membrane potential, firing rates (86, 87), ion conductance, and intracellular Ca^2+^ levels (86, 88, 89). GC signaling decreases glutamatergic inputs and increases GABAergic inputs to postsynaptic peptidergic neurons in the paraventricular nucleus (PVN) of the hypothalamus. Such decrease and increase of glutamatergic and GABAergic inputs are mediated by retrograde release of endocannabinoid and neuronal nitric oxide, respectively (90–92). Some of these findings have been replicated in animal models, yet it needs to be investigated what the neuronal changes observed mean in the context of stress response. Our assay system and what we are learning from modeling stress in zebrafish will be a useful platform to screen for such neural pathways and molecular components.

## Materials and Methods

### Materials and equipment

All the materials and equipment used for this study are listed in the Supplemental Table 1.

### Zebrafish husbandry

Wild-type zebrafish (*Danio rerio*) were purchased from Segrest Farm and maintained in the Zebrafish Core Facility. Fish are handled following standard practices (93) and guidelines from the Institutional Animal Care and Use Committee (IACUC) in the Mayo Clinic (A34513-13-R16, A8815-15). Zebrafish are kept in a 9 L (25–35 adults) or 3 L housing tanks (10–15 adults) at 28.5°C with a light/dark cycle of 14h/10h. Larval zebrafish are raised on petri dish until 5 dpf at which time they were placed in the nursery system. Larval fish are fed on paramecium from 5 to 14 dpf and brine shrimp is introduced on 10 dpf.

### Production and specification of custom light boxes

Light boxes are designed to provide illumination from the bottom (Mayo Clinic Division of Engineering). Light sources are LED diodes on a strip with white or infrared light emission. Light is diffused through white acrylic board to minimize reflection and maximize recording efficiency from a video camcorder (HDR-CX560V, Sony Corp.) mounted at the top of the assay chamber. The dimension of a light box is 18.25” x 20.625”. Multiple units of light boxes were purchased from Super Bright LEDs Inc. and engineered to have dual light sources and a control panel to adjust the intensity outside the box. White light illumination was adjusted to High (dual light assays) or Medium (NaCl/cinnamon oil assays) (Fig. S1). The distribution of wavelength of white light is between 430 and 710 nm. The distribution of wavelength of infrared light was between 790 and 880 nm.

### Locomotor behavioral assays: Light

To obtain embryos, zebrafish adult pairs were placed in a crossing tank that has an inner tank with slits of openings and a divider between a male and female fish a day before spawning. On the following day, embryos are spawned in the morning when the divider is lifted and fallen into the space between the inner and outer tanks. Collected embryos are placed in 100x15 mm petri dishes and kept at 28.5°C in a light/dark cycle of 14h/10h. Unfertilized embryos are eliminated on the same day (0 dpf) and any morphologically abnormal embryos the following day (1 dpf). Fresh 0.5x embryo media is provided (1 dpf) (93). A single larva is placed into each well of a 48-well plate with 500 µL of embryo media on 3 dpf and continuously kept at 28.5°C in a light/dark cycle of 14h/10h in an incubator located in a behavioral assay room to minimize handling of the plates on the assay day.

On 5 dpf, light assays are performed. Larval zebrafish (5 dpf) are acclimated in a pair of 48-well plates in IR (850 nm) for 30 min, exposed to a brief illumination in white light (50 sec), and then back in IR for 20 min (Fig. 1*A*). Their locomotor activity (total distance moved) in each period is video-recorded in IR for 15 min (baseline), white light for 50 sec (treatment), and IR for 20 min (post-treatment). Video recordings (30 fps) in infrared light are performed in the “Nightshot” mode in the video camcorder without using its own infrared beam. Video recordings (30 fps) during white light treatment are performed with a regular recording mode. The difference in locomotion after abrupt light change (IR-white-IR) during the post-treatment period (IR) is quantified with an in-house developed locomotor tracking software (40). The output of the software is a CSV (comma separated values) file with locomotor response information, which was analyzed using R or IBM^®^ SPSS^®^.

### Locomotor behavioral assays: sodium chloride and cinnamon oil

All preparation protocols are the same as the description in light assays. Larval zebrafish (5 dpf) are acclimated in a pair of 48-well plates (450 µL embryo media/well) in white light for 30 min and challenged with 100 mM sodium chloride (Fig. 1*B*). Their locomotion is video-recorded for each period as baseline (15 min) and post-treatment (30 min) before and after sodium chloride (or cinnamon oil) application (50 µL of working stock). Since the purpose of cinnamon oil assay is to show that fish have locomotor response capacity, the initial 10 minutes are used for statistical analysis of cinnamon oil control assays. The final concentration of NaCl is 100 mM, and 7.41 µg/mL (~50 µM) for cinnamon oil. A matching vehicle treatment is used: embryo media for NaCl and DMSO (0.1%) for cinnamon oil.

### Statistical analysis

All data are reported as means ± 95% confidence interval (CI) unless otherwise stated. Statistical analysis is performed using R language (94) and IBM^®^ SPSS^®^ Version 22. For multiple statistical comparisons among several treatment conditions, two-way or three-way mixed analysis of variance (mixed ANOVA) was used. In two-way mixed ANOVA, there were a between-subjects factor and a within-subjects factor. For example, in dual-light assays, the genotype (WT, het, or hom) was a between-subjects factor and the two time-points of locomotor response measurement (labeled as exposure), obtained before or after the white light illumination exposure, were a within-subjects factor. In three-way mixed ANOVA, there were two between-subjects factors and a within-subjects factor. In hyperosmotic stress (NaCl) assays, the genotype (WT, het, or hom) and the treatment condition (VEH *v*. NaCl) were two between-subjects factors. The two time-points of locomotor response measurement (labeled as exposure) obtained before or after the chemical treatment exposure were a within-subjects factor. ANOVA analyses were followed by post-hoc analysis (Tukey’s honest significant difference test). For statistical comparisons involving two treatment groups, the Student’s *t* test was used.

## ACKNOWLEDGMENTS

We thank for their advice on experimental designs: Thesis advisory committee (Stephen C. Ekker, Ph.D., John R. Henley, Ph.D., Joseph A. Murray, M.D., Robin Patel, M.D., Susannah J. Tye, Ph.D.) for Han Lee, zebrafish model systems seminar group (Christopher K. Pierret, Ph.D., Lisa A. Schimmenti, M.D., Caroline R. Sussman, Ph.D., Shizhen (Jane) Zhu, M.D., Ph.D.), and the Neurobiology of Disease program (Allan J. Bieber, Ph.D., DooSup Choi, Ph.D., John D. Fryer, Ph.D., John R. Henley, Ph.D., Pamela J. McLean, Ph.D., Owen A. Ross, Ph.D., Isobel A. Scarisbrick, Ph.D.). For their work in zebrafish husbandry: Mayo Clinic Zebrafish Core Facility (Tammy M. Greenwood, M.S., M.B.A., Danielle E. Hunter, Devin B. Copley, Casey M. Phillips), Mayo Clinic Animal Models and Genetics Core (Camden L. Daby, Melissa S. McNulty, M.S.). For engineering custom light boxes: Mayo Clinic Division of Engineering (Allen K. Rech, PMP, Shaun A. Herring, Roger J. Mahon, Daniel J. (Clay) Mangiameli). For reviewing the manuscript: Louis El Khoury, Ph.D., Noriko Ichino, Ph.D., and Jennifer M. Reiman, Ph.D.

---

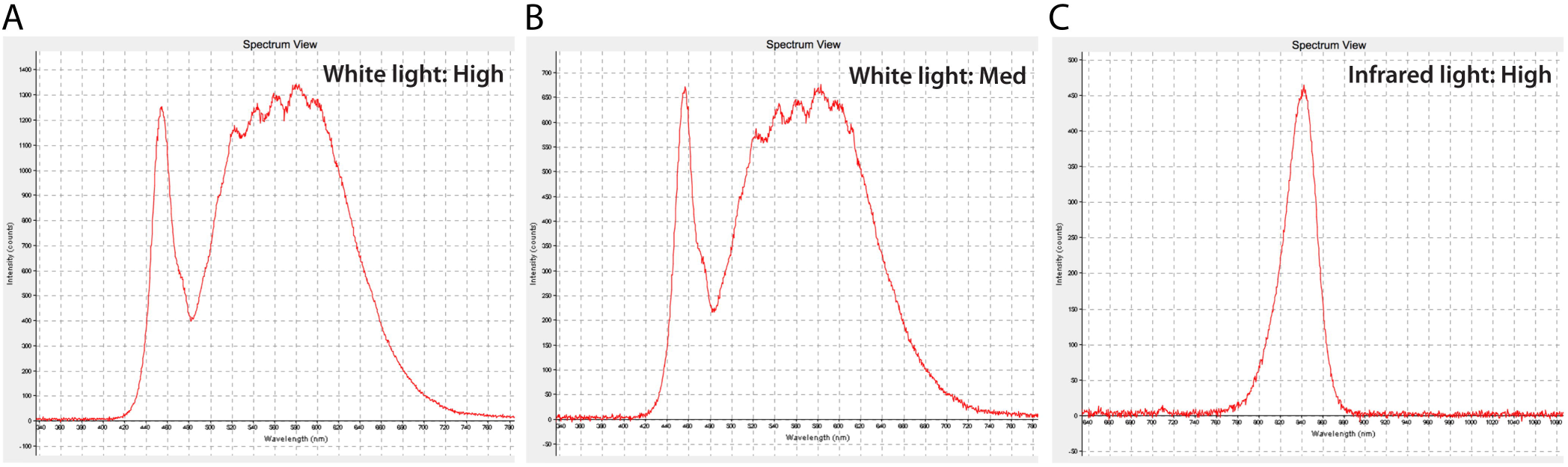

